# ELECTOR: Evaluator for long reads correction methods

**DOI:** 10.1101/512889

**Authors:** Camille Marchet, Pierre Morisse, Lolita Lecompte, Arnaud Lefebvre, Thierry Lecroq, Pierre Peterlongo, Antoine Limasset

## Abstract

The error rates of third-generation sequencing data have been capped above 5%, mainly containing insertions and deletions. Thereby, an increasing number of diverse long reads correction methods have been proposed. The quality of the correction has huge impacts on downstream processes. Therefore, developing methods allowing to evaluate error correction tools with precise and reliable statistics is a crucial need. These evaluation methods rely on costly alignments to evaluate the quality of the corrected reads. Thus, key features must allow the fast comparison of different tools, and scale to the increasing length of the long reads. Our tool, ELECTOR, evaluates long reads correction and is directly compatible with a wide range of error correction tools. As it is based on multiple sequence alignment, we introduce a new algorithmic strategy for alignment segmentation, which enables us to scale to large instances using reasonable resources. To our knowledge, we provide the unique method that allows producing reproducible correction benchmarks on the latest ultra-long reads (longer than 100k bases). It is also faster than the current state-of-the-art on other datasets and provides a wider set of metrics to assess the read quality improvement after correction. ELECTOR is available on GitHub (https://github.com/kamimrcht/ELECTOR) and Bioconda.

## INTRODUCTION

### Motivations

Pacific Biosciences (PB) and Oxford Nanopore Technologies (ONT) long reads, despite their high error rates and complex error profiles, were rapidly adopted for various applications. An increasing number of projects, especially for assembly (1), long-distance haplotyping or structural variant calling (2), indeed benefits from the long-range information these reads provide. These reads display high error rates (from 5%-12%, according to technologies and libraries, to as much as 30% for the oldest datasets), that largely surpass those of Illumina reads. Given these high error rates, the first step of many applications is error correction. However, this stage can be a time bottleneck (2).

Moreover, contrary to Illumina, where the majority of errors are substitutions, long reads mainly contain insertions and deletions (indels) (ONT reads are more deletion-prone, whereas PB reads contain more insertions). This combination of issues requires novel and specific algorithmic developments. To this extent, dozens of error correction methods directly targeting these long reads emerged in the last five years. The first range of error correction tools, called “*hybrid correctors*”, uses both short and long reads to perform error correction, relying on the deep coverage and low error rate of the short reads to enhance long reads sequences. The second group of methods, called “*self-correctors*”, intends to correct long reads with the sole information contained in their sequences (see (3) for a review of correctors). Both paradigms include quite diverse algorithmic solutions, which makes it difficult to globally compare the correction results (in terms of corrected bases, quality, and performances) without a proper benchmark. Besides, the quality of the error correction has considerable impacts on downstream processes. Hence, it is interesting to know beforehand which corrector is best suited for a particular dataset (according to its coverage, its error rate, the sequenced genome, or the sequencing technology, for instance). Developing methods allowing to evaluate error correction tools with precise and reliable statistics is, therefore, a crucial need.

Methods for evaluating correctors should allow tracking the novelties of the methods. Indeed, since long read technologies still evolve, current correctors implementations are prone to many changes. Methods for evaluating correctors must be usable on datasets of various complexity (from bacteria to eukaryotes) to reproduce a wide variety of possible scenarios. They also should be fast and lightweight, and should not be orders of magnitude more resource and time consuming than the actual correction methods they evaluate. This aspect is particularly critical, since correction evaluators also stand in the perspective of new correction methods developments, as they can help to provide accurate and quick comparisons to the state-of-the-art. For developers as well as users, correction evaluators should describe with precision the correction method’s behavior (i.e., the number of corrected bases, introduced errors or read breakups) to identify its potential pitfalls.

### Previous works

Works introducing novel correction methods usually evaluate the quality of their tools based on how well the corrected long reads realign to the reference.

Despite being useful, this information remains incomplete. In particular, it is likely not to mention poor quality reads, or regions to which it is difficult to align.

In (6), La *et al.* introduced a new way to obtain metrics describing the quality of the error correction that does not solely present the similarities between the aligned corrected reads and the reference genome. Relying on simulated data, they proposed the idea of a three-way alignment between the reference genome, the uncorrected reads, and the corrected reads. They presented results on PB data for hybrid error correction tools, by introducing LRCstats, an evaluation tool aiming at answering to the problematics above.

With its three-way alignment scheme, LRCstats provides reads’ error rate before and after correction, as well as the detailed count of every type of error. However, only studying the reads’ error rate after correction is not a satisfying indication of the corrector’s behavior. For instance, there is no clue about the putative insertions of new errors by the corrector. To perform such analysis of the method’s pros and cons, we need additional metrics such as precision (relevant corrected bases among all bases modified by the corrector) and recall (correct bases that have been retrieved by the corrector among all bases to be corrected). Such metrics have already been proposed in earlier works dedicated to short reads, such as the error correction evaluation toolkit introduced in (7). However, this contribution is out of the scope of this work. Indeed, algorithms to process short reads are different from those at stake in our case, due to the length, the high error rates, and the complex error profiles of the long reads.

Moreover, LRCstats relies on a multiple alignment scheme which suffers from high resource needs when processing large numbers of reads, i.e., when coverage or genome sizes are large. For the same reason, LRCstats alignment scheme becomes limited when sequences to process grow. However, the sequencing depth and the length of the long reads keep on increasing, especially with so-called ONT ultra-long reads (up to 1M bases) starting to appear in recent works for larger genomes (8). Moreover, deep coverages are expected to help the correction of very long sequences (2). Thus, novel methods must be proposed in order to evaluate the correction of such datasets in a reasonable amount of time.

### Contribution

To cope with the identified limits of LRCstats, we propose ELECTOR, a new evaluation tool for long read error correction methods. ELECTOR can evaluate the correction of simulated as well as real long read datasets, provided a reference genome is available for the sequenced species. It takes as input a reference genome in FASTA format, a set of corrected reads in FASTA format, and the corresponding uncorrected reads, either via a FASTA format file in the case of real data or via the suite of files provided by the simulator in case of simulated data. In its output, ELECTOR provides a broader range of metrics than LRCstats, which evaluates the actual quality of the correction. In particular, it measures recall, precision, and error rate for each read. ELECTOR also informs about typical difficulties long read correctors can encounter, such as homopolymers, and reads that have been trimmed, split or extended during the correction. Finally, it also provides reads remapping and assembly metrics.

In order for ELECTOR to provide these additional metrics, we propose a novel multiple sequence alignment (MSA) strategy. This new algorithmic approach is designed to allow the MSA computation to scale to ultra-long reads and to large datasets of several billions of base-pairs. It compares in a fast way three different versions of each read: the *corrected* version, the *uncorrected* version and the *reference* version, which is a substring of the reference genome. For each read, we perform a MSA of its triplet. A key idea of this strategy is a divide-and-conquer approach that divides the reads into smaller sequences with an anchoring process, and thus allows to compute several smaller MSAs. These multiple, smaller MSAs, are then combined to obtain the final MSA, of the whole length of the sequences. The anchoring process is designed to work with erroneous sequences and takes into account gapped alignment due to truncated corrected reads. We believe that the interests of this novelty are not limited to the ELECTOR framework. Indeed, it may be a useful strategy for any domain requiring MSAs of long and highly erroneous sequences.

For simulated reads, ELECTOR is compatible with state-of-the-art long reads simulation tools, such as NanoSim (9) or SimLoRD (10), on which introduced errors are precisely known. Moreover, it is meant to be a user-friendly tool, that delivers its results through different output formats, such as graphics that can be directly integrated into the users’ projects. This tool was designed to be directly compatible with a wide range of state-of-the-art error correction tools without requiring any pre-processing by the user. In particular, ELECTOR is compatible with the latest self-correction methods, and we thus present novel results on such tools, that were not tackled by LRCstats.

## MATERIAL AND METHODS

### Input sequences

ELECTOR is compatible with long reads simulators SimLoRD and NanoSim, and real read sequences (see Figure 1 for an overview of ELECTOR). When using long reads simulated with one of these tools, the reference sequences are directly retrieved by ELECTOR, by parsing the files generated during the simulation. When using these state-of-the-art long reads simulation tools, we ensure to take as input sequences that closely simulate the actual characteristics of the long reads. However, it is possible to use other long reads simulation tools. In this case, the user must provide the *reference* sequences itself. The genome used for the simulation, the files generated by the simulator, and the corrected reads, output by the desired correction method, are then provided as an input. ELECTOR then compares three different versions of each read: the *uncorrected* version, as provided by the sequencing experiment or by the read simulator, the *corrected* version, as provided by the error correction method, and the *reference* version, which is a portion of the reference genome, representing a perfect version of the original read, without any error. For real data, the *reference* sequences are retrieved by aligning the *uncorrected* reads to the reference genome, using Minimap2. Only the best hit for each read is kept and used to determine the corresponding *reference* sequence. In the case a read cannot align to the reference genome, and thus cannot produce a *reference* sequence, the read is excluded from the analysis.

**Figure 1.**
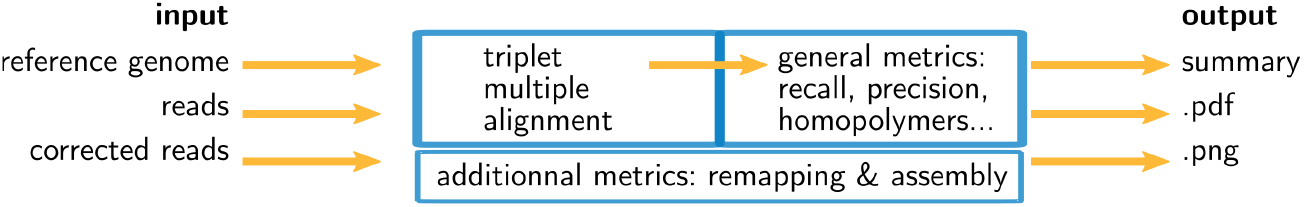
Overview of ELECTOR. Inputs are the sequences at different stages: without errors (from the reference genome), with errors (simulated or real uncorrected reads) and corrected (after running a correction method). We compute a multiple sequence alignment of the three versions of each sequence and analyze the results to provide correction quality measures. In order to provide additional information, reads are assembled using Minimap2 (4) and Miniasm (5) and both the reads and the contigs are aligned to the reference genome. A text summary, plots, and a pdf summary are output.

### Scalable triplet multiple alignment

With real or simulated reads, the core of the algorithmic novelty is to propose the comparison of the three different versions of each read (*reference*, *uncorrected*, and *corrected*) in a triplet multiple alignment. These three versions of each read undergo a multiple sequence alignment, to collect their differences and similarities at each position of the alignment.

*Principle* With the three versions of each read, our triplet multiple alignment strategy computes a MSA, using a partial order alignment algorithm. The MSA is initialized with the *reference* sequence, and the *corrected* and *uncorrected* sequences are then sequentially added. This step yields a multiple alignment matrix that is output in pseudo FASTA (PIR) format for each triplet. The triplet multiple alignments are computed using an implementation of partial order alignment graphs (11). Partial order alignment graphs are used as structures containing the information of the multiple aligned sequences. To this extent, a directed acyclic graph (DAG) contains the previous multiple sequence alignment result. The vertices store consecutive nucleotides from the sequences. Each new sequence is aligned to this DAG in a generalization of the Needleman-Wunsch algorithm. Paths in the DAG represent the successive alignments.

However, such a procedure can be time-consuming when applied to noisy long reads (see Table 2). Thus, we propose a novel multiple sequence alignment heuristic. We recall the values of all the parameters mentioned in the following paragraphs in Supplementary Table S1.

#### Segmentation strategy for the MSA

To reduce the computation time of our approach, we propose a segmentation strategy, as sketched in Figure 2. It consists of dividing the global multiple alignment into several smaller instances. Drawing inspiration from MUMmer’s (12) and Minimap’s (5) longest increasing subsequence approaches, we select a sequence of seeds *S*_1_,*…S*_*N*_ that can be found (in the given order) within the three sequences. From this sequence of seeds, we extract the *N* +1 substrings (*W*_0_,*W*_1_,*…,W*_*N*_) delimited by the seeds in the three versions of the read. We thus extract *str*[0: *position_S*_1_],*str*[*position_S*_1_ + *length_S*_1_: *position_S*_2_],*…, str*[*position_S*_*i*_ +*length_S*_*i*_: *position_S*_*i*+1_],*…, str*[*position_S*_*N*_: *str_size*], with *str* being the sequence of the *reference*, *corrected* or *uncorrected* version, and call these substrings *windows*. We, therefore, compute independent MSAs for each window triplet (*W*_*i*__*reference,W*_*i*__*corrected,W*_*i*__*uncorrected*), and then reconstitute the global multiple alignment by concatenation. We now describe the procedure more in detail. For each triplet, we compute the *k*-mers that will be used as seeds (called *seed k*-mers) so that they comply with the following properties:

1. They appear in each of the three versions of the sequence.
2. They are not repeated across any of the versions of the sequence.
3. They are not overlapping in any of the versions of the sequence.

**Figure 2.**
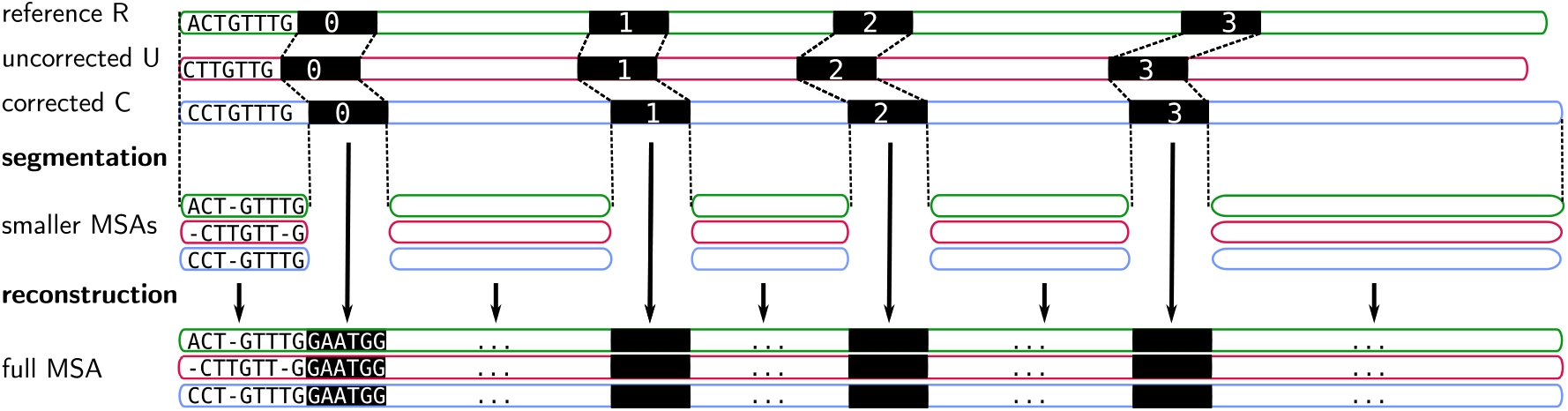
Segmentation strategy to compute a multiple sequence alignment for a triplet of *reference*, *uncorrected* and *corrected* versions of a read. Instead of computing a MSA on the whole length of the sequences, we rather divide this problem into smaller instances. As each version is different, to decide where to start and end the alignments, we look for seed *k*-mers (in black) that are exact local matches between the three sequences. We then compute individual, separate MSAs, for subsequences bordered by seeds (or located at the extremities of the sequences). These multiple MSAs are finally concatenated, along with the seed *k*-mers, to obtain a single, full MSA, of the whole length of the sequences.

Using dynamic programming, the longest increasing subsequence of seed *k*-mers, *S*_1_,*…S*_*N*_, is computed. Pairs of successive seed *k*-mers, *S*_*i*_,*S*_*i*+1_ delineate windows. The size of these seed *k*-mers is adapted according to the current observed error rates (5, 13), and ranges between 9 and 15 nucleotides. As it is difficult to *a priori* select a *k*-mer size, we designed a quick iterative strategy that tries several values of *k*, to choose the most suitable for a given triplet. Starting from *k* = *k*_*max*_ (set to 15 by default), we keep on decreasing *k* until the size of the largest window no longer decreases. Whenever the largest window’s size no longer decreases, or *k*_*min*_ (set to 9 by default) is reached, the process stops. Minimizing the size of the largest window as such allows us to ensure that we compute MSAs on the smallest possible windows, in order to reduce the computational costs as much as possible.

Once windows are computed, we produce MSAs of each window triplet (*W*_*i*__*reference,W*_*i*__*corrected,W*_*i*__*uncorrected*) independently, as described in the previous paragraph, using subsequently smaller alignment graphs. Finally, the multiple small MSAs are concatenated, along with the seed *k*-mers, to obtain a single MSA of the whole length of the read triplet.

If we were able to bound the size of the windows, we could guarantee an asymptotic time linear to the read length for the alignment computation. In practice, our implementation can produce large windows, but we observe a running time almost linear in the length of the reads, as shown in our experimental results.

To avoid computing metrics on poorly corrected reads, we filter out corrected reads length is below a given parameter (see Supplementary Table S1 for its default value) and triplets for which no seed *k*-mers can be found. These two types of filtered reads are tagged and reported apart in ELECTOR’s outputs to inform the user about their numbers.

#### Handle reads of different sizes in the segmentation strategy

In the case of a truncated corrected read (trimmed/split), the *corrected* version is shortened in comparison to the two other versions. A part of the *reference* and *uncorrected* sequences is thus missing in the *corrected* sequence. A prefix, a suffix, or both, can be missing depending on the case. Trimmed and split scenarios are outlined in Figure 4. As we only use anchors shared among the three sequences, in the case of a missing prefix in the corrected version, *W*_0__*reference* and *W*_0__*uncorrected* will, therefore, be larger than *W*_0__*corrected* (see an example of a missing suffix in Figure 3). Computing a MSA between those three sequences would thus be irrelevant. Furthermore computing a MSA on two possibly long sequences (as a large sequence may be missing) is pricey. As corrected reads can be truncated at the beginning, at the end, or both, the symmetrical scenario can occur for suffixes.

**Figure 3.**
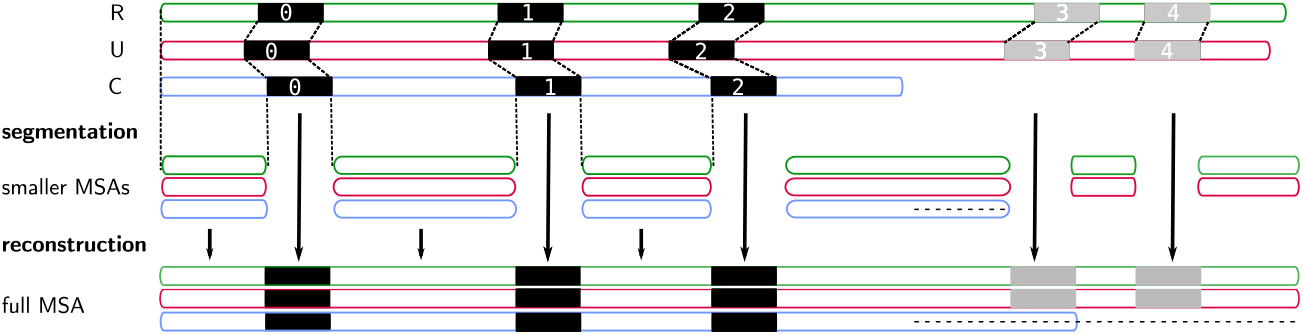
Segmentation strategy when the *corrected* read is smaller. As in Figure 2, R,U,C stand for the reference, uncorrected and corrected read triplet. Here, the *corrected* read is shortened on its right end. To avoid passing subsequences starting from seed 2 to the end of each sequence to the MSA, which would be costly to compute, we perform a second segmentation strategy. This strategy allows us to retrieve a new set of seeds (gray seeds 3 and 4). This new set of seeds divides the remaining subsequences (suffixes in this case) in *reference* and *uncorrected* into windows on which we compute MSA separately. The full MSA is reconstructed by concatenation, and dots are added to complete the *corrected* MSA line.

**Figure 4.**
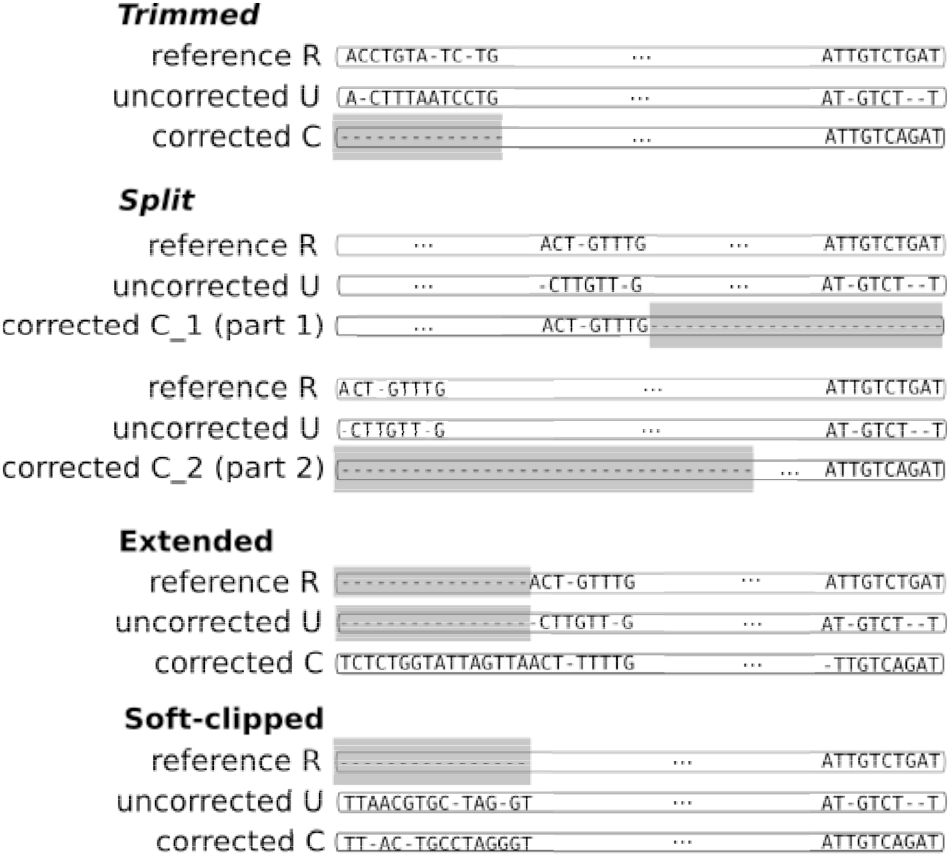
Three scenarios of corrected read categories in MSA results. Trimmed reads have a *corrected* version with a missing prefix and/or suffix (grey region). Split reads have been fragmented into several parts during the correction, and subsequences can be missing between consecutive fragments (grey regions). Extended corrected reads have a *corrected* version with an additional prefix and/or suffix which is (are) not present in the two other versions (missing regions in grey). Soft-clipped reads have a *reference* version with a missing prefix and/or suffix (grey region).

To cope with this problem, we detect such cases by checking the length of the first windows. If *W*_0__*reference* and *W*_0__*uncorrected* are large (1000 nucleotide) and at least 2 time larger than *W*_0__*corrected*, we use a segmentation scheme only with *k*-mers from *reference* and *uncorrected*, and only align their two prefixes.

This way, we can efficiently compute a MSA when the corrected reads do not cover the whole genome region they originally come from, avoiding to run a MSA on large/unrelated sequences. The procedure is symmetrical for suffixes.

This procedure is essential for correctors that output numerous split reads, which would induce extremely long run-time due to large sequence MSA computations described before.

### Inference of quality evaluation metrics from MSA

#### Classification of corrected reads

ELECTOR reports different categories of *corrected* reads: regular reads, trimmed/split reads, extended reads, soft-clipped reads, bad quality reads and short reads. Figure 4 shows how we deduce the trimmed, split and extended categories from the MSA result.

**Regular** reads are neither trimmed, split, extended nor soft-clipped.

**Trimmed** reads are reads that lack a part of their prefix, suffix, or both (first scenario in Figure 4).

**Split** reads are reads composed of several fragments that come from a single original read, that could only be corrected on several distinct parts (second scenario in Figure 4). Split reads are aligned as trimmed reads are. However, in the case of split reads, we gather all fragments that come from a single initial read, in order to build a single MSA from the several, distinct MSAs induced by the different fragments. Supplementary Figure S1 illustrates this process.

We thus report how many reads were trimmed or split during the correction. Moreover, for each trimmed or split corrected read, we report the total uncorrected length of its associated *reference* read (i.e., the length that is not covered by any fragment).

**Extended** reads are reads that have a prefix and/or a suffix that was not present in the *reference* sequence (third scenario in Figure 4). These reads can be chimeras from the correction step, and can, for instance, come from chimeric connections between unrelated parts of the graph (14) or the assembly of unrelated short reads (15).

However, they can also be reads that were over-corrected by a graph-based correction method, that kept on traversing the graph after reaching the *uncorrected* reads’ extremities. We do not compute quality evaluation metrics on the extended regions, but we report the number of extended reads, as well as their mean extension size, with respect to the *reference* reads.

We define a split/trimmed/extended region as the prefix or suffix (or both) of the MSA in which no *corrected* nucleotide appears (for split and trimmed), or no *uncorrected* and *reference* nucleotide appear (for extended). These regions are represented in grey in Figure 4.

**Soft-clipped** reads are reads for which the extremities were soft clipped during the alignment to the reference genome (last scenario in Figure 4). This category only arises in real data mode, as we only retrieve *reference* reads by aligning the *uncorrected* reads to the reference genome in this case. For such reads, we do not compute quality evaluation metrics on the soft clipped regions, as they could not be appropriately aligned to the reference genome, and were therefore not used to determine the *reference* read.

**Bad quality** reads are low-quality reads that were removed before the MSA step, to avoid computing metrics on poorly corrected reads. As mentioned previously, these are the reads for which no seed *k*-mers were found during the segmentation process. These reads are tagged and reported apart in ELECTOR’s output, to inform the user about their number. We only report their number as no metric can be computed since they are not aligned.

**Short reads** are reads which are shorter than ℓ% of the *reference* sequence length (ℓ being a parameter set to 10 by default). As for the bad quality reads, these reads are also removed before the MSA step, and only the number of such reads is reported.

#### Recall, precision, error rate

Once the MSA is computed, we have access to information about the differences and similarities in nucleotide content for each position of the three versions of a sequence. Insertions and deletions are represented by a “.” in the deleted parts, and by the corresponding nucleotide (`A`,`C`,`T` or `G`) in the inserted parts. Let us denote, respectively, by *nt*(*R,p*_*i*_),*nt*(*C,p*_*i*_),*nt*(*U,p*_*i*_) the characters of *reference, corrected* and *uncorrected* versions in {*A,C,G,T,.*}, at position *p*_*i*_ (0 ≤ *i≤ N*), in a MSA of size *N*. Figure 5 shows how recall and precision are computed. The set 𝒫 of positions to correct is composed of positions *p*_*i*_ such as *nt*(*R,p*_*i*_) ≠ *nt*(*U,p*_*i*_). The set *ε* of existing positions in the corrected version is defined by including any position *p*_*x*_ from the *corrected* version that is not counted in a trimmed/split/extended/soft-clipped region. The processed positions set 𝒞 is defined as 𝒫∪{*p*_*j*_ /*nt*(*C,p*_*j*_) ≠ *nt*(*R,p*_*j*_)} ∩𝓔. The correct positions set 𝒞*o* is defined as 𝒞∩{*p*_*j*_ /*nt*(*C,p*_*j*_)= *nt*(*R,p*_*j*_)}. The recall, precision and error rate are computed as follows:

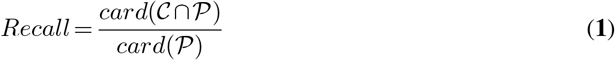

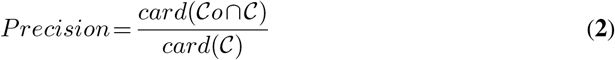

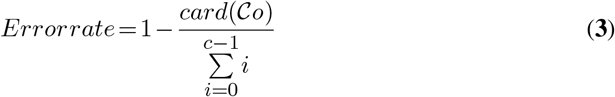

with *c* the length of the corrected read.

**Figure 5.**
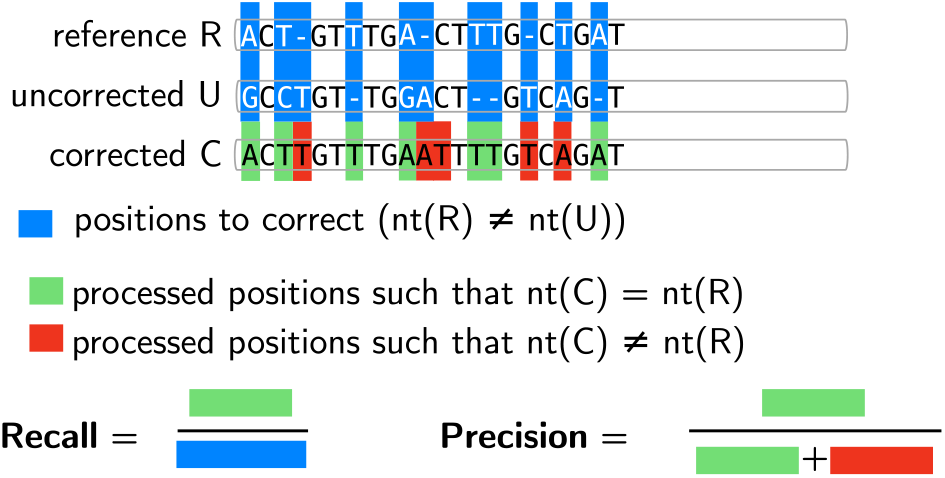
Computation of recall and precision using triple base-wise comparison at each MSA’s position. *nt*(*R*) (respectively *nt*(*U*),*nt*(*C*)) represents the character in *reference* (respectively *uncorrected, corrected*) line of the MSA at a given position.

#### Additional metrics

ELECTOR provides the number of trimmed or split corrected reads, and the mean missing size of these reads, as well as the number of extended reads, and the mean extension size of these reads. The size distribution of the sequences, before and after correction, is reported graphically.

In the case of split reads, we report the length of each fragment in the distribution. The %GC of the *corrected* and *reference* reads is also output, as well as the total number of insertions, deletion, and substitution, in the *uncorrected* and *corrected* reads. ONT reads are known to be more error-prone than PB reads in homopolymers. Thus, we propose metrics to examine these particular regions We show the ratio of homopolymer sizes in the *corrected* version over the *reference* version. The closer it is to one, the better the corrector overcame possible systematic errors in ONT reads.

More details on the computation of the insertions, deletions, substitutions counts, and on the ratio of homopolymer sizes are shown, respectively, in Supplementary Figure S2 and S3.

#### Remapping of corrected reads

In addition to all previously presented metric computations, we also take advantage of the presence of the reference genome to evaluate corrected reads quality. We perform remapping of the corrected reads to the reference genome using Minimap2. We report the number of corrected reads, the total number of bases, the average length of the reads, the percentage of aligned reads, the mean identity of the alignments, as well as the genome coverage, i.e., the percentage of bases of the reference genome to which at least a nucleotide aligned.

#### Post-correction assembly metrics

Again, in addition to metrics obtained thanks to our MSA strategy, we assess the correction quality through its consequences on the assembly quality of the corrected reads. We perform the assembly of the corrected reads using Miniasm (5), as we mainly seek to develop a pipeline providing fast results. We acknowledge that assemblers such as Smartdenovo (16) or Canu (17) are more sensitive, but as they display much larger runtimes, Miniasm provides a satisfying compromise.

As for the metrics of the assembly, we output the overall number of contigs, the number of contigs that could be aligned, the number of breakpoints of the aligned contigs, the NGA50 and NGA75 sizes of the aligned contigs, as well as the genome coverage. Using the assemblies that we provide, further analyses can be performed using dedicated software such as QUAST-LG (18).

We also perform the alignment of the contigs with Minimap2. The computation of the different metrics, for remapping and assembly assessment, is then performed by parsing the generated SAM file.

## RESULTS

### Validation of the segmentation strategy for MSA

To validate our segmentation strategy for MSA, we show to which extent its results differ from the classical MSA approach. In particular, we expect that recall, precision, and error rate hardly differs, thus showing that both behaviors produce very similar results. Conversely, we expect a decisive gain in time with our segmentation strategy compared to the original algorithm. We thus compared multiple alignment results obtained with our strategy to results obtained with the regular implementation of partial order alignment graphs on multiple datasets of different read lengths, which affects the run-time of the alignments. Results are presented in Table 2. We observe that while the two strategies provide very similar metrics, the segmentation strategy can reduce the runtime by orders of magnitude compared to the regular approach, especially when the reads grow longer.

### Validation on synthetic datasets

In this section, we present the results of ELECTOR and LRCstats on several simulated datasets from different species. Further details about these datasets are given in Table 1. The choice of synthetic data was motivated by the need to know the *reference* sequences (which are portions of the reference genome, representing perfect versions of the original reads, on which no error would have been introduced) to precisely control the results brought by the assessed correction method.

**Table 1.** Description of the datasets used in our experiments. ^1^https://www.ncbi.nlm.nih.gov/nuccore/NC_000913 ^2^www.genoscope.cns.fr/externe/nas/references/yeast/W303_pacbio_assembly.fa.gz ^3^https://www.ebi.ac.uk/ena/data/view/GCA_000002985.3 ^4^Only chromosome 1 was used. https://www.ncbi.nlm.nih.gov/nuccore/NC_000001.11 ^5^Only reads from chromosome 1 were used.

| Dataset | *E. coli* | *S. cerevisiae* | *C. elegans* | *H. sapiens* |
| --- | --- | --- | --- | --- |
| **Reference organism** |  |  |  |  |
| Strain | K-12 substr. MG1655 | W303 | Bristol N2 | GRCh38 |
| Reference sequence | NC_000913<sup>1</sup> | scf718000000{084-13}<sup>2</sup> | GCA_000002985.3<sup>3</sup> | NC_000001.11<sup>4</sup> |
| Genome size | 4.6 Mbp | 12.2 Mbp | 100 Mbp | 249 Mbp |
| **Simulated Pacific Biosciences data** |  |  |  |  |
| Number of reads | 11,306 | 30,132 | 244,277 | - |
| Average length (bases) | 8,226 | 8,204 | 8,204 | - |
| Number of bases (M bases) | 93 | 247 | 2,004 | - |
| Coverage | 20x | 20x | 20x | - |
| Error rate (%) | 18.6 | 18.6 | 18.6 | - |
| **Real Oxford Nanopore data** |  |  |  |  |
| Accession | - | - | - | PRJEB23027<sup>5</sup> |
| Number of reads | - | - | - | 1,075,867 |
| Average length (bases) | - | - | - | 6,744 |
| Number of bases (M bases) | - | - | - | 7,256 |
| Coverage | - | - | - | 29x |
| Error rate (%) | - | - | - | 17.60 |

**Table 2.** Comparison of the two multiple alignment strategies on simulated *E. coli* datasets. Three datasets were simulated, with a 10% error rate, a coverage of 100x, and a fixed read length of 1k bases, 10k bases and 100k bases, respectively. The reads were corrected using Canu with default parameters.

| Experiment | Recall (%) | Precision (%) | Error rate (%) | Time |
| --- | --- | --- | --- | --- |
| "1k" MSA | 99.712 | 98.996 | 1.02 | 2h 05 min |
| "1k" segmentation +MSA | 99.769 | 98.992 | 1.021 | **28 min** |
| "10k" MSA | 99.921 | 99.781 | 0.206 | 20 h 50 min |
| "10k" segmentation + MSA | 99.921 | 99.795 | 0.207 | **29 min** |
| "100k" MSA | 99.913 | 99.925 | 0.044 | 8 days 18 h 38 min |
| "100k" segmentation +MSA | 99.924 | 99.903 | 0.098 | **1 h 11 min** |

#### ELECTOR sample output

As previously mentioned, ELECTOR computes general metrics: recall, precision, error rate, among other metrics, and provides a graphic representation of their distributions.

A subset of the metrics produced by ELECTOR using reads corrected by the following tools: HALC (19), HG-CoLoR (20), LoRDEC (21), Canu (17), Daccord (Tischler, G., & Myers, E. W. (2017). Non hybrid long read consensus using local de Bruijn graph assembly. bioRxiv, 106252.) and MECAT (22) is presented in Table 3. These metrics are consistent with the results presented in the respective tools’ publications. The whole set of metrics, including remapping and assembly assessment, are presented in Supplementary Table S3 and S4.

**Table 3.**
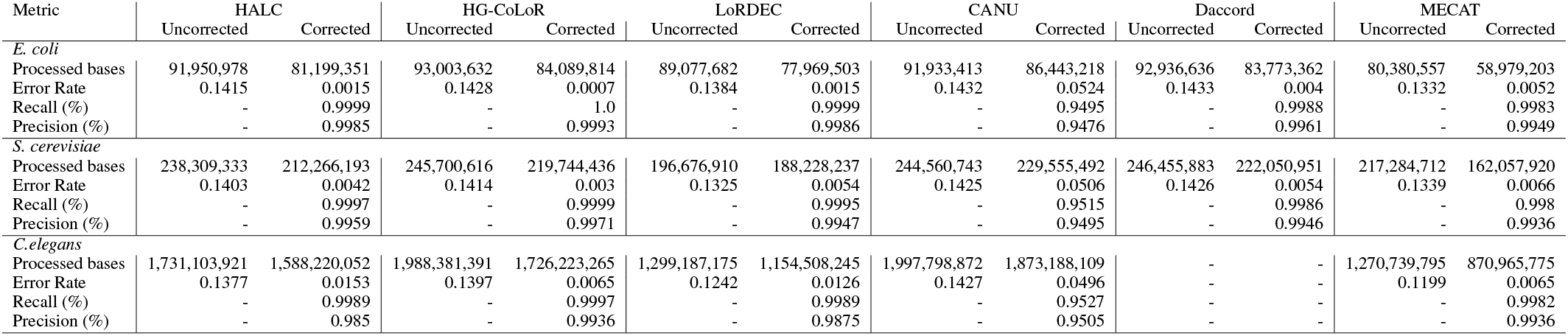
Examples of the main metrics reported by ELECTOR on *E. coli, S. Cerevisiae* and *C. elegans* datasets. A dash in the Uncorrected columns indicates that the metric is not computed for the *uncorrected* reads. Daccord could not be run on the *C. elegans* dataset, and reported an error.

#### Comparison to state-of-the-art

Recently, several benchmark analysis were proposed for long reads (comparison of hybrid correction methods (23), comparison of hybrid and self-correction methods (Zhang, H. et al. (2019). A comprehensive evaluation of long read error correction methods. BioRxiv, 519330.), analysis of long read correction on transcriptomic reads (24)). In this work, we focus on the methodological basis allowing to efficiently perform and reproduce such benchmarks, rather than highlighting the pros and cons of available correction methods. The presented correction performances are thus showed for validation purposes and are not intended to be a benchmark of existing correction methods. In the rest of the result section, we report comparisons to the only other automated evaluation tool for long reads correction: LRCstats.

In Table 4, we compare the metrics displayed by ELECTOR and LRCstats. Correction of the *S. cerevisiae* dataset by HALC (a hybrid correction method) and Canu (self-correction method), are evaluated and reported as an example output. The complete results provided by LCRstats and ELECTOR, for each correction tool, and on each dataset, are presented in Supplementary Tables S2 and S3.

**Table 4.**
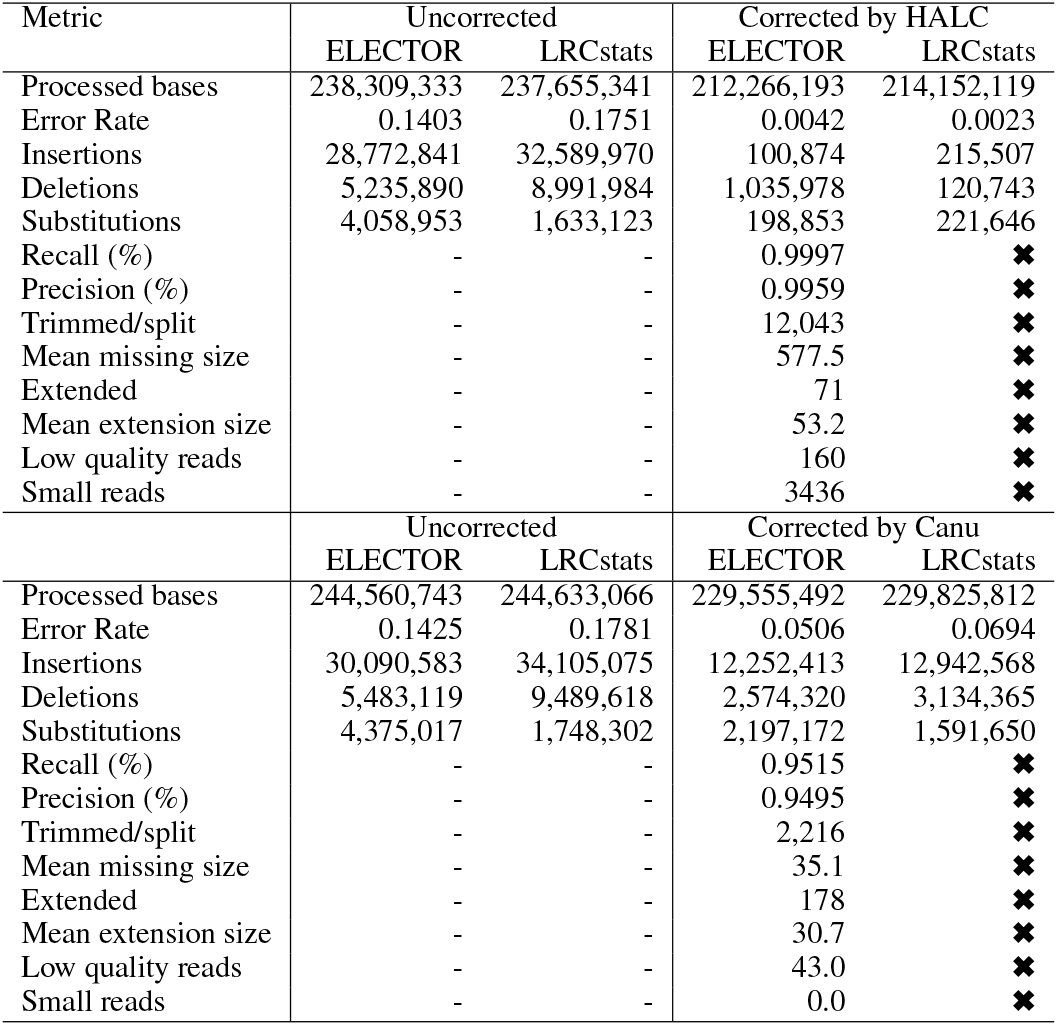
Comparison of ELECTOR’s and LRCstats’s outputs. Both tools were evaluated on the *S. cerevisiae* dataset, using a hybrid corrector (HALC) and a self corrector (Canu). A dash in the Uncorrected columns indicates that the metric is not computed for the *uncorrected* reads. A cross indicates that LRCstats does not provide the metric.

Both LRCstats and ELECTOR compute metrics on *corrected* reads and the corresponding *uncorrected* sequences of those reads (reported respectively as *corrected* and *uncorrected*).

The first result to notice in Table 4 is that the error rates and the amount of processed bases announced in the *uncorrected* reads can differ from one correction method to the other, both for ELECTOR and LRCstats. Such differences can be explained by the fact that HALC and Canu do not correct the same set of reads, which leads to different set of *uncorrected* reads to evaluate.

As ELECTOR and LRCstats rely on different rules to exclude reads from the analysis, and do not align split reads in the same way, we observe that they do not process the same quantity of reads.

LRCstats concatenates the different parts of a split read before aligning the concatenation, even if a missing region can exist between two consecutive fragments. This behavior can complicate the alignment task and introduce a bias in the output metrics. On the contrary, ELECTOR processes the different fragments separately before reconstituting the whole alignment and thus takes into account missing regions. These differences thus have an impact on the metrics displayed for corrected reads. ELECTOR processes slightly more bases than LRCstats on the two studied datasets. However, reads falling into particular categories (very short reads and low-quality reads) are not taken into account in ELECTOR’s counts, and are reported apart, while they are absent from LRCstats’s output.

Different alignment strategies in both tools also have impacts on the results, which explains the differences seen in indels and substitutions counts. However, ELECTOR and LRCstats globally report the same trends of two successful corrections that decreased the error rates.

Additional metrics, specific to ELECTOR, point out noteworthy differences between the two correction methods, such as the high quantity of trimmed or split reads when using HALC in comparison to Canu. These metrics are essential for further steps such as assembly since less advantage is taken from shortened reads to resolve repeats. They also help to understand more in-depth the correctors’ behavior. In this example, Canu corrects with lower recall and precision than HALC, but this is nuanced because ELECTOR reports it produces less trimmed/split reads.

### Performance comparison

In this section, we compare LRCstats and ELECTOR runtime and memory consumption on several datasets chosen to represent different use cases. Results are presented in Table 5 and 6. For the experiments presented in Table 5, both tools were ran on a 20-core cluster node equipped with 250 GB of RAM. For the experiments presented in Table 6, we used a 16-core computer equipped with 64 GB of RAM. In order to compare similar operations, ELECTOR’s runtime and memory consumption do not consider the remapping and assembly steps. We present the metrics and resource consumption of this module apart, in Supplementary Table S4.

**Table 5.** Evolution of ELECTOR and LRCstats runtime and memory consumption according to the read length. The datasets were simulated from the *E. coli* genome, with a 100x coverage, a 10% error rate, and fixed read length of 1k bases, 10k bases, 100k bases, and 1M bases. The reads were corrected by Canu, using default parameters.

| Tool | Read length (bases) | Memory (MB) | Elapsed time | CPU time |
| --- | --- | --- | --- | --- |
| LRCstats | 1k | 1,803 | 42 min | 8 h 18 min |
| ELECTOR | 1k | 1,030 | 12 min | 28 min |
| LRCstats | 10k | 13,484 | 4 h 51 min | 70 h 38 min |
| ELECTOR | 10k | 3,091 | 13 min | 29 min |
| LRCstats | 100k | - | - | - |
| ELECTOR | 100k | 12,231 | 28 min | 1 h 11 min |
| LRCstats | 1M | - | - | - |
| ELECTOR | 1M | 24,881 | 2 h 44 min | 11 h 05 min |

**Table 6.**
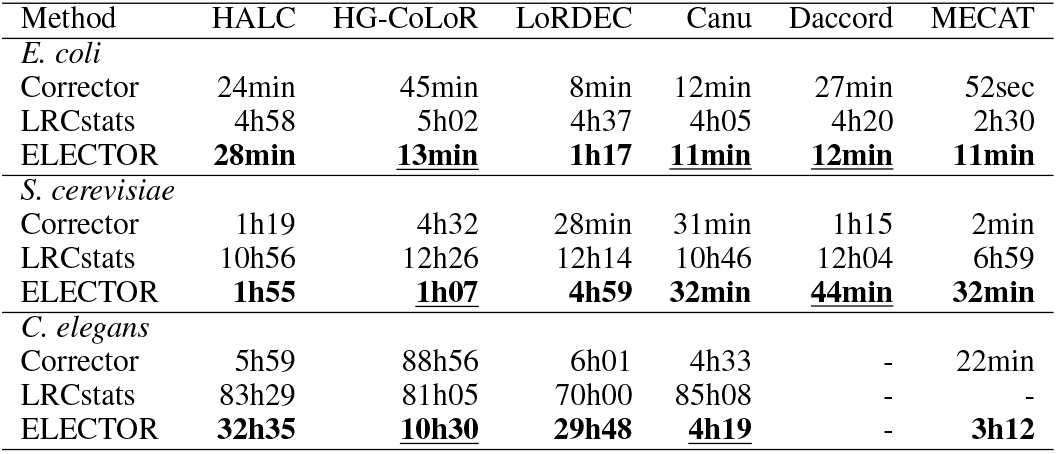
Runtimes of ELECTOR and LRCstats on different datasets and different correction tools. Both ELECTOR and LRCstats were launched with 9 threads. The different correction methods were launched with 16 threads. The runtimes of the correctors are also included as a matter of comparison. The fastest evaluation method is shown in bold for each case. When the evaluation method is also quicker than the correction method itself, it is underlined. Daccord could not be run on the *C. elegans* dataset, and reported an error. LRCstats crashed while assessing the *C. elegans* dataset corrected by MECAT.

| Method | HALC | HG-CoLoR | LoRDEC | Canu | Daccord | MECAT |
| --- | --- | --- | --- | --- | --- | --- |
| *E. coli* |  |  |  |  |  |  |
| Corrector | 24min | 45min | 8min | 12min | 27min | 52sec |
| LRCstats | 4h58 | 5h02 | 4h37 | 4h05 | 4h20 | 2h30 |
| ELECTOR | **28min** | **13min** | **1h17** | **11min** | **12min** | **11min** |
| *S. cerevisiae* |  |  |  |  |  |  |
| Corrector | 1h19 | 4h32 | 28min | 31min | 1h15 | 2min |
| LRCstats | 10h56 | 12h26 | 12h14 | 10h46 | 12h04 | 6h59 |
| ELECTOR | **1h55** | **1h07** | **4h59** | **32min** | **44min** | **32min** |
| *C. elegans* |  |  |  |  |  |  |
| Corrector | 5h59 | 88h56 | 6h01 | 4h33 | - | 22min |
| LRCstats | 83h29 | 81h05 | 70h00 | 85h08 | - | - |
| ELECTOR | **32h35** | **10h30** | **29h48** | **4h19** | - | **3h12** |

We first assess, in Table 5, the performances of both tools on several simulated *E. coli* datasets with different read lengths, ranging from 1k bases to 1M bases. As expected, the runtime and memory consumption of both tools grow with the read length. However, ELECTOR can handle reads larger than 10k bases better than LRCstats, thanks to its segmentation strategy. In particular, ELECTOR is several orders of magnitude faster than LRCstats on the 10k bases experiment, and can also handle longer reads, up to 1M bases, using moderate resources. LRCstats was much more memory consuming and was thus unable to run on reads longer than 10k bases, despite having access to 250 GB of RAM. These results underline that ELECTOR can scale to extremely long reads. Considering the ever-growing length of the long reads and the tremendous impact of such very long sequences, we believe that this ability is a significant advantage of ELECTOR obtained thanks to its segmentation technique.

We also observe, in Supplementary Table S5, that the error rate of the input reads has a negligible impact on the performances of the tools.

In Table 6, we compare the performances of different correctors with the time needed to evaluate their outputs, using ELECTOR and LRCstats. Interestingly, we observe that LRCstats is mostly slower than the correction step itself, which is not desirable. ELECTOR is often faster than or comparable to the corrector itself, except for MECAT which is distinctly efficient. These reduced runtimes could be a beneficial gain for benchmark analysis, and could also be critical for the development of new correction methods. Another observation from Table 6 is that we can notice large divergence in ELECTOR runtimes on the same dataset corrected by different tools. This behavior can be due to two factors. On the one hand, ELECTOR’s runtime optimization is prone to be more or less pronounced according to the read length (segmentation is expected to be easier with larger reads) and quality of the correction (more errors make it more difficult to find common seeds). On the other, ELECTOR’s runtime is also related to the number of split corrected reads output by the corrector. Indeed, a larger number of split reads implies a more significant number of triplet multiple alignments, and thus an increased runtime. In particular, in the experiments presented here, the largest runtimes can be observed for the evaluation of LoRDEC and HALC on the *C. elegans* dataset. As shown in Supplementary Table S3, these tools are also those that produced the most considerable amount of trimmed/split reads on this dataset. A way to accelerate ELECTOR analysis would be to adapt its parameters to avoid small read fragments.

### Simulations for the validation of ELECTOR’s real data mode

In order to validate ELECTOR’s real data mode, we ran the following experiment. We used a simulated dataset, and we assessed its correction using the two different modes of ELECTOR: simulated and real data. First, we ran it classically, by providing the simulation files as an input, so that ELECTOR could retrieve the actual *reference* reads by parsing the files. Second, we ran it by only providing the FASTA file of simulated reads as an input, so ELECTOR had to retrieve the *reference* reads by aligning the uncorrected long reads to the reference genome, as if they were not simulated. We ran this experiment on the *S. cerevisiae* dataset. To further validate ELECTOR’s behavior on real data, we assessed the correction of both a hybrid corrector, HALC, and a self-corrector, Canu. Results of these experiments are shown in Table 7.

**Table 7.** Comparison of the results output by ELECTOR, using simulated and real data modes. The two experiments were run on the same *S. cerevisiae* dataset, using a hybrid corrector (HALC) and a self corrector (Canu).

| Metric | Uncorrected |  | Corrected by Halc |  |
| --- | --- | --- | --- | --- |
|  | Simulated | Real | Simulated | Real |
| Processed bases | 238,309,333 | 238,119,170 | 212,266,193 | 212,141,319 |
| Error Rate | 0.1403 | 0.1449 | 0.0042 | 0.0104 |
| Recall (%) | - | - | 0.9997 | 0.9938 |
| Precision (%) | - | - | 0.9959 | 0.9897 |
| Insertions | 28,772,841 | 26,796,500 | 100,874 | 90,737 |
| Deletions | 5,235,890 | 5,042,365 | 1,035,978 | 1,490,680 |
| Substitutions | 4,058,953 | 3,682,863 | 198,853 | 182,590 |
| Trimmed/split | - | - | 12,043 | 13,320 |
| Mean missing size | - | - | 577.5 | 896.0 |
| Extended | - | - | 71.0 | 39.0 |
| Mean extension size | - | - | 53.2 | 72.0 |
| Low quality reads | - | - | 160.0 | 152.0 |
| Small reads | - | - | 3436.0 | 3438.0 |

| Metric | Uncorrected |  | Corrected by Canu |  |
| --- | --- | --- | --- | --- |
|  | Simulated | Real | Simulated | Real |
| Processed bases | 244,560,743 | 244,402,568 | 229,555,492 | 229,403,697 |
| Error Rate | 0.1425 | 0.1442 | 0.0506 | 0.052 |
| Recall (%) | - | - | 0.9515 | 0.9499 |
| Precision (%) | - | - | 0.9495 | 0.9481 |
| Insertions | 30,090,583 | 28,452,967 | 12,252,413 | 10,965,458 |
| Deletions | 5,483,119 | 5,800,286 | 2,574,320 | 2,916,564 |
| Substitutions | 4,375,017 | 4,081,445 | 2,197,172 | 1,940,888 |
| Trimmed/split | - | - | 2216.0 | 4943.0 |
| Mean missing size | - | - | 35.1 | 74.7 |
| Extended | - | - | 178.0 | 169.0 |
| Mean extension size | - | - | 30.7 | 31.9 |
| Low quality reads | - | - | 43.0 | 42.0 |
| Small reads | - | - | 0.0 | 0.0 |

We observe that ELECTOR’s results are consistent, both in simulated and real data mode. In particular, recall and precision are very similar. The two modes display some differences in the input uncorrected reads (as shown by the amount of processed bases), that have an impact on the differences observed between their results.This behavior is due to the bias induced by the additional alignment step that the real data mode requires. The main differences that appear occur on metrics that are highly dependent on the alignment results, such as the number of trimmed, split and extended reads, and the sizes of these events; as well as indels and substitutions counts.

### Results on a real human dataset

To demonstrate ELECTOR’s results in a realistic scenario for large genomes, we evaluate the correction of a real human dataset. We report results, as well as runtime of the evaluation, in Table 8. The reads were corrected with MECAT, using default parameters, before running ELECTOR. Using 20 threads, we were able to obtain the results for the 650,771 corrected reads of the dataset in less than 19 hours. Results reported by ELECTOR show that MECAT can correct human reads with a 20% error rate with more than 90% of recall and precision, which is consistent with the published results.

**Table 8.** Evaluation of the correction of a real human dataset with ELECTOR. The reads were corrected with MECAT, using default parameters, before the evaluation. ELECTOR evaluated a total of 650,771 reads. Small reads are corrected reads which length is lower than 10% of the original read. Low quality corrected reads are reads for which an insufficient number of seeds was found during the segmentation process. Homopolymer ratio is the ratio of homopolymer sizes in *corrected* vs *reference*. We reported the wallclock time of the run, using 20 threads.

|  | Uncorrected | Corrected with MECAT |
| --- | --- | --- |
| Processed bases | 5,605,157,590 | 5,451,767,836 |
| Recall(%) | - | 92.70 |
| Precision(%) | - | 91.50 |
| Error rate | 0.1974 | 0.0861 |
| Trimmed/split | - | 570,635 |
| Mean missing size | - | 362.0 |
| Extended | - | 275 |
| Mean extension size | - | 62.4 |
| Low quality reads | - | 4,279 |
| Small reads | - | 356 |
| Insertions | 247,953,086 | 10,144,736 |
| Deletions | 746,165,024 | 473,239,036 |
| Substitutions | 162,822,923 | 7,521,389 |
| Homopolymer ratio | - | 0.7570 |
| Runtime | - | 18 h 27 min |

## DISCUSSION

In ELECTOR, we propose a novel efficient algorithmic approach of segmentation strategy for multiple sequence alignment. We adapted this task for this original and specific application of long reads comparison. New segmentation strategies for MSA were recently proposed (Nogales, E. G. et al. (2018). Fast and accurate large multiple sequence alignments using root-to-leave regressive computation. bioRxiv, 490235.). However, these methods are not specifically designed for noisy long reads. On such data, both the high error rates and lengths are troublesome factors for the multiple sequence alignment computation. In such a perspective, a generalization of our segmentation strategy, allowing long reads multiple sequence alignments of more than three sequences would be very interesting. Such a generalization could indeed be relevant for critical applications such as assembly, consensus or variant detection.

ELECTOR’s real data mode uses a prior alignment of the reads to a reference genome, in order to retrieve the *reference* versions of the reads. We demonstrated that ELECTOR’s metrics in its real data mode remain highly similar to what would be obtained in its simulated mode. However, we can point out two limitations of ELECTOR. First, even if the data can come from an actual sequencing experiment, a reference genome needs to exist for the sequenced species, in order to retrieve the *reference* reads, and thus perform the evaluation. Second, we encourage users to be very cautious about ELECTOR’s results on real data, especially when looking at the number of trimmed, split, or extended reads and at the sizes of such events. Indeed, these metrics are highly dependent on the result of the alignment of the *uncorrected* reads to the reference. These metrics can thus be subject to errors, especially when aligning relatively short or highly erroneous/chimeric reads, or reads coming from repeated regions.

A future application is the evaluation of correction methods directly targeted at RNA long reads sequencing. As shown in a recent study (24), RNA long reads have specific requirements that are not met by current methods, which calls for new correctors in the future. ELECTOR could be coupled with a reference transcriptome or a RNA long read simulator, although, currently, such a simulation software does not exist to our knowledge.

## CONCLUSION

We presented ELECTOR, a tool that enables the evaluation of self and hybrid long reads correction methods, and that allows evaluating the behavior of a given correction tool in a controlled situation. ELECTOR provides a wide range of metrics that include base-wise computation of recall, precision, error rates of corrected and uncorrected reads as well as insertions, deletions and substitutions counts, and homopolymers correction. In particular, we believe that recall and precision are of prime interest to characterize a correction tool behavior. Indeed, these metrics allows spotting specific pitfalls, or undesired effects, that remain unclear when only looking at the error rates of the corrected reads. ELECTOR reports a text summary of its different metrics, along with pdf and png versions, including plots of the key figures. This allows users to easily integrate ELECTOR’s outputs into reports.

Even though ELECTOR relies on multiple sequence alignment techniques that can be very resource-consuming, we were able to evaluate the behavior of a representative list of state-of-the-art hybrid and self-correctors, ran on reads from small bacterial to large mammal genomes. We also showed that ELECTOR’s performances allow it to scale to very long reads, displaying lengths up to 1M bases, with moderate resource needs.

In particular, ELECTOR is typically faster than most error correction methods. ELECTOR’s ability to quickly handle real-world datasets with low memory consumption is preeminently valuable when working on long read exploitation routines, and represents a significant improvement in comparison to the state-of-the-art.

The efficiency of ELECTOR relies on an innovative and promising segmentation algorithm for multiple sequence alignment of noisy long reads. This procedure drastically reduces the time footprint of the multiple sequence alignment, making it able to scale to very long sequences. We believe this algorithm could be improved and applied to a broad range of applications implying multiple sequence alignment of long, noisy sequences.

## ACKNOWLEDGEMENTS

We thank Pierre Marijon for his help with the Bioconda integration. Part of this work was performed using the computing resources of CRIANN (Normandy, France).

## SUPPLEMENTARY MATERIAL

### Dealing with split corrected sequences

In order to compute recall, precision and correct base rates only on actually corrected parts, even if the correction did produce some split reads, we designed a particular procedure (Figure 6).

**Figure 6.**
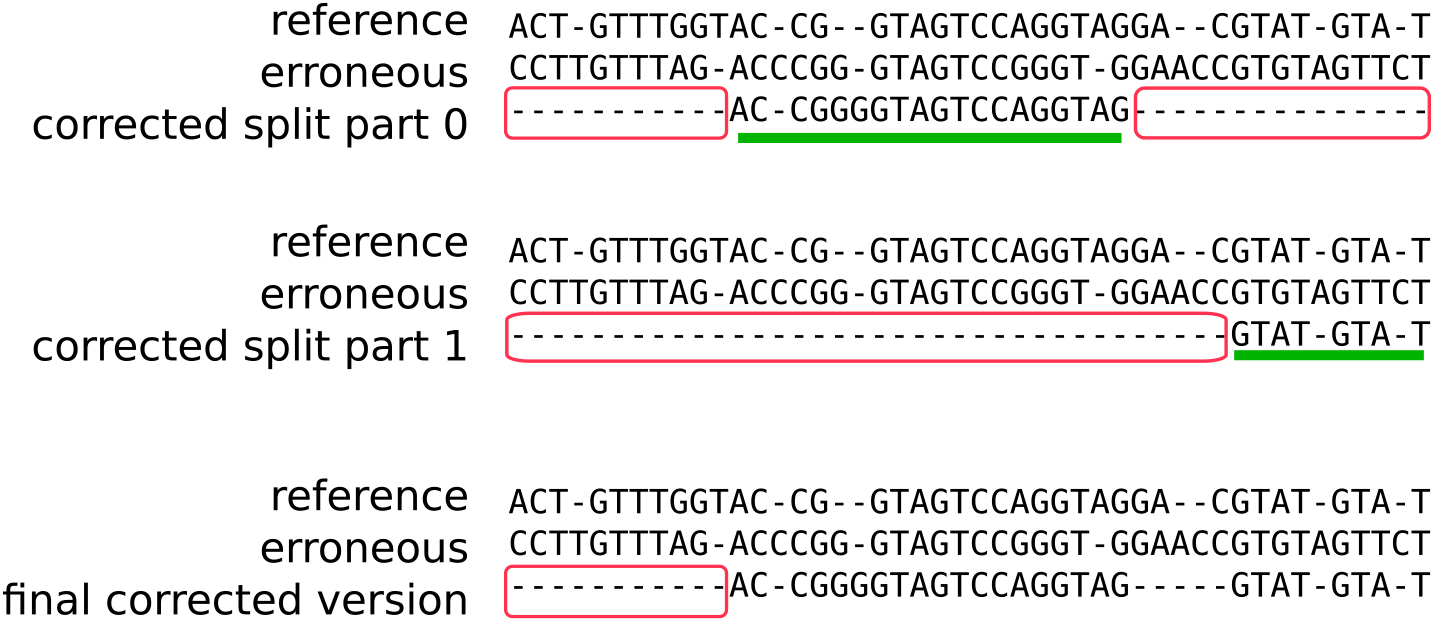
In this example, the read is corrected in two distinct parts. After *reference* and *uncorrected* sequences duplication, we have two distinct triplets. Large gaps in the MSA (represented in red boxes), which lengths are over a given threshold, are identified. Only large gaps are considered as non-corrected regions, in order to conserve smaller gaps that correspond to indels. Remaining regions (underlined in green) are considered as corrected. Then, the different corrected regions are combined into a single MSA (bottom of the figure), and large gaps are identified once again. Recall, precision and correct base rate are computed on the final corrected zone. The missing size in the *corrected* read is computed by counting the number of nucleotides in the *reference* read that are included in the red window.

### Metrics details

In Figures 7,8 we give further details on how metrics about insertions, deletions and substitutions counts, as well as homopolymers, are computed.

**Figure 7.**
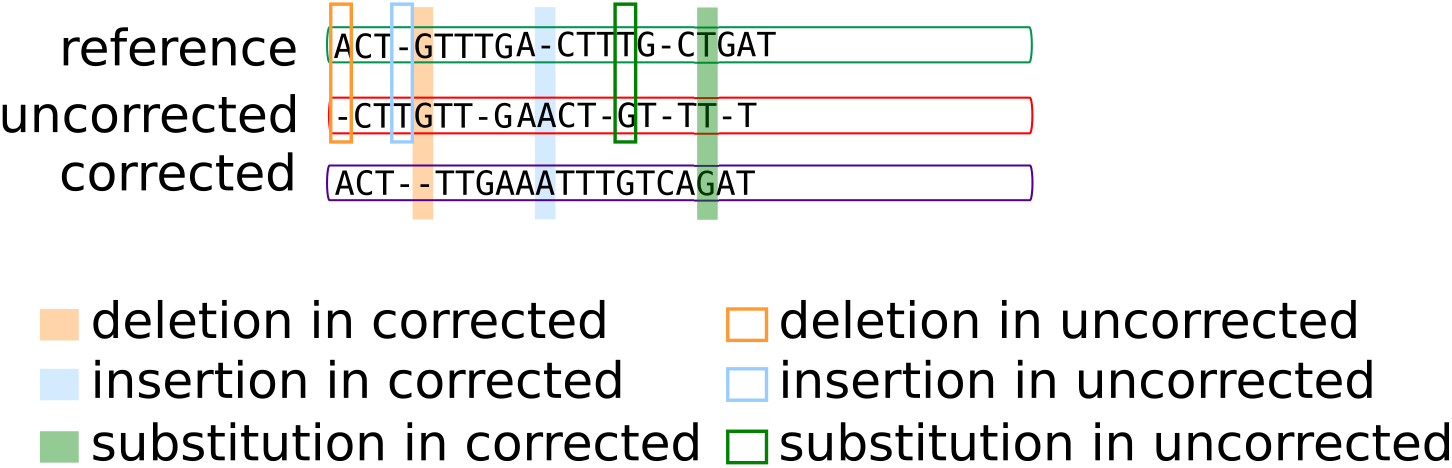
In this example, we show how the numbers of insertions, deletions, and substitutions are computed in the *uncorrected* and *corrected* reads. For each scenario, one example is pictured. For instance, A deletion in the corrected read is shown in filled orange, the reference nucleotide is G and the uncorrected nucleotide is G. Both the *corrected* and the *uncorrected* versions are compared base-to-base to the *reference* version in order to obtain these counts. Insertions, deletions, and substitutions are not computed in parts of the MSA that correspond to extended, trimmed, split, or soft-clipped regions.

**Figure 8.**
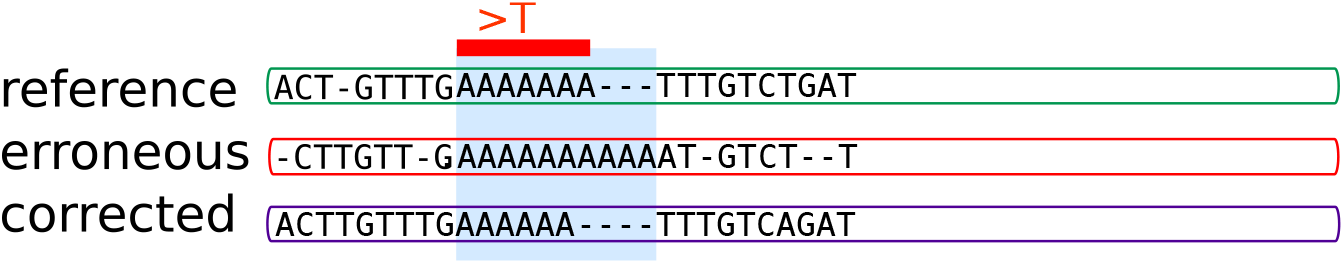
Homopolymers are detected as mono-nucleic chains longer than a given threshold (6 in practice) in the *reference* read. In ELECTOR, we display the ratio of the homopolymers size in the *corrected* reads over the *reference* reads. In this example, the ratio would be 6/7.

**Table 9.** ELECTOR’s default parameters. ^1^ the alignment matrix coefficients are the same as those used in LRCstats.

|  |  |
| --- | --- |
| *k* size (max/min) | 15/9 |
| alignment scores (opening/insertion in gap) | 10/5<sup>1</sup> |
| min % of reference read size under which a corrected read is filtered out | 10 |
| min number of seeds under which a corrected read is filtered out | 1 |
| min size of the first/last window to start the segmented gap strategy | 1000 |
| min gap size to consider a sequence as split, trimmed or extended | 20 |
| min homopolymer size | 5 |

**Table 10.**
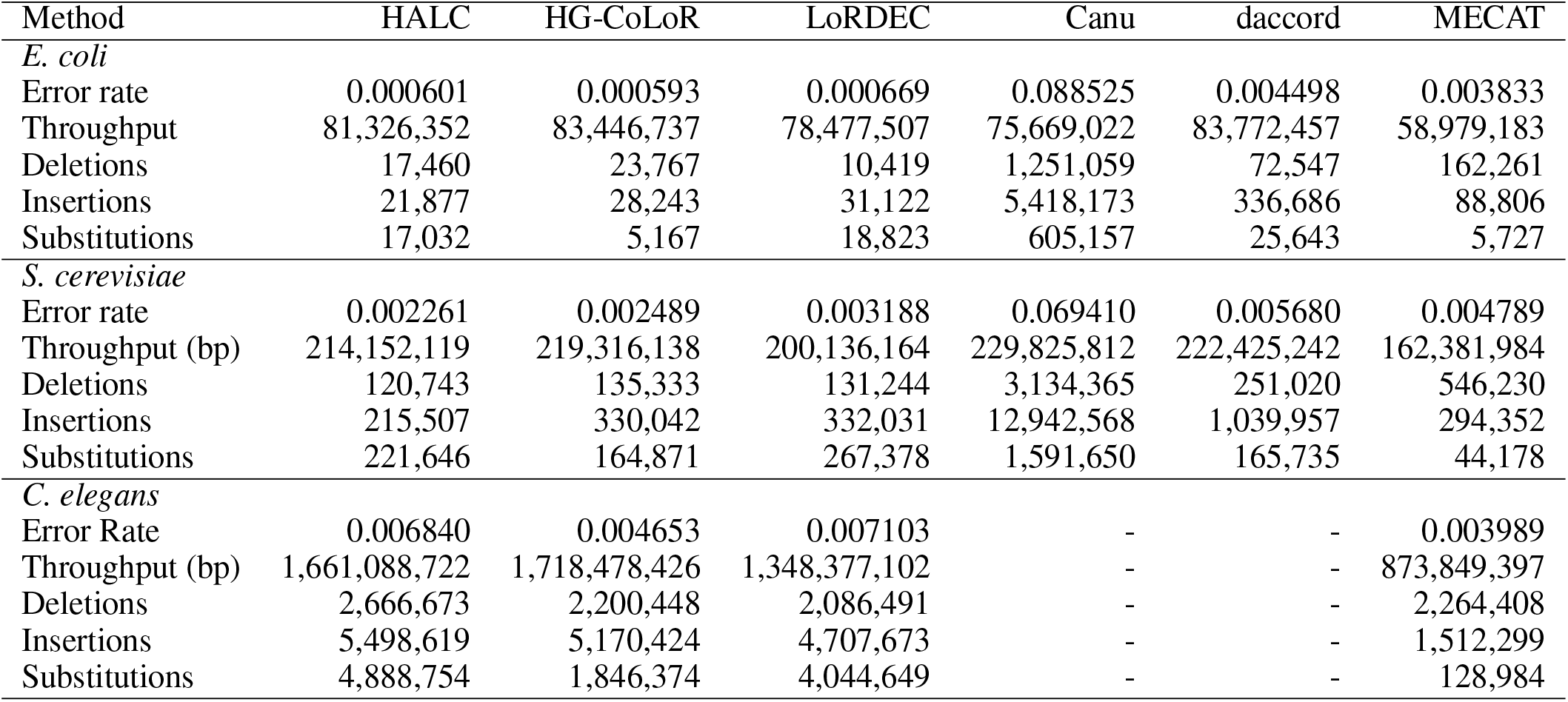
Statistics of the long reads after correction with the different methods, as reported by LRCstats. Daccord could not be run on the *C. elegans* dataset, and reported an error, and LRCstats crashed on the *C. elegans* dataset corrected by Canu.

**Table 11.** Statistics of the long reads after correction with the different methods, as reported by ELECTOR. Daccord could not be run on the *C. elegans* dataset, and reported an error.

| Method | HALC | HG-CoLoR | LoRDEC | Canu | daccord | MECAT |
| --- | --- | --- | --- | --- | --- | --- |
| *E. coli* |  |  |  |  |  |  |
| Throughput (bp) | 81,199,351 | 84,089,814 | 77,969,503 | 86,443,218 | 83,773,362 | 58,979,203 |
| Error Rate | 0.0015 | 0.0007 | 0.0015 | 0.0524 | 0.0040 | 0.0052 |
| Recall (%) | 0.9999 | 1.0000 | 0.9999 | 0.9495 | 0.9988 | 0.9983 |
| Precision (%) | 0.9985 | 0.9993 | 0.9986 | 0.9476 | 0.9961 | 0.9949 |
| Insertions | 15,034 | 33,409 | 21,205 | 4,804,729 | 321,532 | 82,121 |
| Deletions | 115,973 | 28,955 | 128,344 | 1,011,985 | 75,627 | 267,547 |
| Substitutions | 13,764 | 3,147 | 17,896 | 841,679 | 33,860 | 7,575 |
| Trimmed/split | 3,816 | 498 | 4,786 | 764 | 119 | 6,304 |
| Mean missing size | 547.3 | 274.5 | 1,033.9 | 27.2 | 369.9 | 2,114.5 |
| Extended | 24 | 8,180 | 0 | 66 | 0 | 0 |
| Mean extension size | 31.1 | 71.1 | 0 | 27.4 | 0 | 0 |
| Low quality reads | 2 | 0 | 0 | 0 | 0 | 0 |
| Small reads | 254 | 4 | 737 | 0 | 4 | 0 |
| *S. cerevisiae* |  |  |  |  |  |  |
| Throughput (bp) | 212,266,193 | 219,744,436 | 188,228,237 | 229,555,492 | 222,050,951 | 162,057,920 |
| Error Rate | 0.0042 | 0.0030 | 0.0054 | 0.0506 | 0.0054 | 0.0066 |
| Recall (%) | 0.9997 | 0.9999 | 0.9995 | 0.9515 | 0.9986 | 0.9980 |
| Precision (%) | 0.9959 | 0.9971 | 0.9947 | 0.9495 | 0.9946 | 0.9936 |
| Insertions | 100,874 | 548,419 | 122,252 | 12,252,413 | 977,499 | 263,724 |
| Deletions | 1,035,978 | 339,174 | 1,385,989 | 2,574,320 | 470,101 | 975,507 |
| Substitutions | 198,853 | 79,127 | 229,925 | 2,197,172 | 200,541 | 52,975 |
| Trimmed/split | 12,043 | 4,562 | 22,470 | 2,216 | 991 | 16,034 |
| Mean missing size | 577.5 | 677.4 | 857.6 | 35.1 | 410.4 | 2,069.5 |
| Extended | 71 | 19,237 | 3 | 178 | 0 | 1 |
| Mean extension size | 53.2 | 72.5 | 1,285.3 | 30.7 | 0 | 23 |
| Low quality reads | 160 | 99 | 252 | 43 | 49 | 50 |
| Small reads | 3,436 | 496 | 22,811 | 0 | 54 | 0 |
| *C. elegans* |  |  |  |  |  |  |
| Throughput (bp) | 1,588,220,052 | 172,6223,265 | 1,154,508,245 | 1,873,188,109 | - | 870,965,775 |
| Error Rate | 0.0153 | 0.0065 | 0.0126 | 0.0496 | - | 0.0065 |
| Recall (%) | 0.9989 | 0.9997 | 0.9989 | 0.9527 | - | 0.9982 |
| Precision (%) | 0.9850 | 0.9936 | 0.9875 | 0.9505 | - | 0.9936 |
| Insertions | 2,520,363 | 5,156,404 | 1,584,355 | 97,517,920 | - | 1,566,201 |
| Deletions | 35,970,722 | 9,986,160 | 23,642,857 | 20,233,048 | - | 5,379,870 |
| Substitutions | 2,919,936 | 775,253 | 1,845,645 | 17,594,481 | - | 139,668 |
| Trimmed/split | 153,855 | 71,079 | 210,051 | 14,407 | - | 120,325 |
| Mean missing size | 777.2 | 866.3 | 1,350.9 | 29.5 | - | 2,320.8 |
| Extended | 772 | 140,791 | 30 | 1,462 | - | 41 |
| Mean extension size | 40.9 | 72.2 | 156.9 | 32.5 | - | 354.7 |
| Low quality reads | 2,582 | 621 | 1,966 | 305 | - | 583 |
| Small reads | 170,934 | 8,849 | 474,725 | 0 | - | 1 |

#### Assembly metrics

In Table 12, we display metrics, runtime and memory consumption for the assessment of the HALC corrected reads on the *S. cerevisiae* dataset.

**Table 12.** Metrics reported by the remapping and assembly module of ELECTOR, on the *S. cerevisiae* dataset, corrected with HALC.

|  |  |
| --- | --- |
| **Remapping** |  |
| Number of reads | 38,926 |
| Number of bases | 214,233,733 |
| Average length | 5,504 |
| Aligned reads (%) | 99.7971 |
| Average identity (%) | 99.4886 |
| Genome coverage (%) | 98.9989 |
| **Assembly** |  |
| Number of contigs | 168 |
| Aligned contigs | 168 |
| Number of breakpoints | 5 |
| NGA50 | 94,358 |
| NGA75 | 55,321 |
| Genome coverage (%) | 91.7579 |
| Time | 3 min |
| Memory (MB) | 2,646 |

**Table 13.** Evolution of ELECTOR’s and LRCstats’ runtime and memory consumption according to the error rate. The datasets were simulated from the *E. coli* genome, with a 50x coverage, a fixed read length of 10 kbp and error rates of 1, 5, 10 and 15%. The reads were corrected by MECAT, using default parameters.

| Tool | Error rate (%) | Memory (MB) | CPU time |
| --- | --- | --- | --- |
| LRCstats | 1 | 13,833 | 147 h 20 min |
| ELECTOR | 1 | 3,899 | 41 min |
| LRCstats | 5 | 13,109 | 142 h 12 min |
| ELECTOR | 5 | 3,433 | 41 min |
| LRCstats | 10 | 12,942 | 140 h 5 min |
| ELECTOR | 10 | 3,097 | 43 min |
| LRCstats | 15 | 12,484 | 122 h 4 min |
| ELECTOR | 15 | 2,837 | 42 min |

## REFERENCES

1. Gordon, D., Huddleston, J., Chaisson, M. J., Hill, C. M., Kronenberg, Z. N., Munson, K. M., Malig, M., Raja, A., Fiddes, I., Hillier, L. W., et al. (2016) Long-read sequence assembly of the gorilla genome. Science, 352(6281), aae0344.

2. Sedlazeck, F. J., Lee, H., Darby, C. A., and Schatz, M. C. (2018) Piercing the dark matter: bioinformatics of long-range sequencing and mapping. Nature Reviews Genetics, p. 1.

3. Laehnemann, D., Borkhardt, A., and McHardy, A. C. (2015) Denoising DNA deep sequencing data-high-throughput sequencing errors and their correction. Briefings in bioinformatics, 17(1), 154–179.

4. Li, H. (2018) Minimap2: pairwise alignment for nucleotide sequences. Bioinformatics, 1, 7.

5. Li, H. (2016) Minimap and miniasm: fast mapping and de novo assembly for noisy long sequences. Bioinformatics, 32(14), 2103–2110.

6. La, S., Haghshenas, E., and Chauve, C. (2017) LRCstats, a tool for evaluating long reads correction methods. Bioinformatics, 33(22), 3652–3654.

7. Yang, X., Chockalingam, S. P., and Aluru, S. (2012) A survey of error-correction methods for next-generation sequencing. Briefings in bioinformatics, 14(1), 56–66.

8. Jain, M., Koren, S., Miga, K. H., Quick, J., Rand, A. C., Sasani, T. A., Tyson, J. R., Beggs, A. D., Dilthey, A. T., Fiddes, I. T., et al. (2018) Nanopore sequencing and assembly of a human genome with ultra-long reads. Nature biotechnology, 36(4), 338.

9. Yang, C., Chu, J., Warren, R. L., and Birol, I. (2017) NanoSim: nanopore sequence read simulator based on statistical characterization. GigaScience, 6(4), 1–6.

10. Stöcker, B. K., Köster, J., and Rahmann, S. (2016) Simlord: Simulation of long read data. Bioinformatics, 32(17), 2704–2706.

11. Lee, C., Grasso, C., and Sharlow, M. F. (2002) Multiple sequence alignment using partial order graphs. Bioinformatics, 18(3), 452–464.

12. Delcher, A., Salzberg, S., and Phillippy, A. (2003) Using MUMmer to identify similar regions in large sequence sets. Current Protocols in Bioinformatics, Chapter 10.

13. Chaisson, M. J. and Tesler, G. (2012) Mapping single molecule sequencing reads using basic local alignment with successive refinement (BLASR): application and theory. BMC bioinformatics, 13(1), 238.

14. Miclotte, G., Heydari, M., Demeester, P., Rombauts, S., Van de Peer, Y., Audenaert, P., and Fostier, J. (2016) Jabba: hybrid error correction for long sequencing reads. Algorithms for Molecular Biology, 11, 10.

15. Madoui, M.-A., Engelen, S., Cruaud, C., Belser, C., Bertrand, L., Alberti, A., Lemainque, A., Wincker, P., and Aury, J.-M. (2015) Genome assembly using Nanopore-guided long and error-free DNA reads. BMC genomics, 16(1), 327.

16. Ruan, J. Smartdenovo: Ultra-fast De Novo Assembler Using Long Noisy Reads. https://github.com/ruanjue/smartdenovo (2017).

17. Koren, S., Walenz, B. P., Berlin, K., Miller, J. R., Bergman, N. H., and Phillippy, A. M. (2017) Canu: scalable and accurate long-read assembly via adaptive k-mer weighting and repeat separation. Genome research, pp. gr–215087.

18. Mikheenko, A., Prjibelski, A., Saveliev, V., Antipov, D., and Gurevich,A. (2018) Versatile genome assembly evaluation with QUAST-LG. Bioinformatics, 34(13), i142–i150.

19. Bao, E. and Lan, L. (2017) HALC: High throughput algorithm for long read error correction. BMC bioinformatics, 18(1), 204.

20. Morisse, P., Lecroq, T., and Lefebvre, A. (2018) Hybrid correction of highly noisy long reads using a variable-order de Bruijn graph. Bioinformatics, 34(24), 4213–4222.

21. Salmela, L. and Rivals, E. (2014) LoRDEC: accurate and efficient long read error correction. Bioinformatics, 30(24), 3506–3514.

22. Xiao, C.-L., Chen, Y., Xie, S.-Q., Chen, K.-N., Wang, Y., Han, Y., Luo, F., and Xie, Z. (2017) MECAT: fast mapping, error correction, and de novo assembly for single-molecule sequencing reads. Nature Methods, 14(11), 1072.

23. Fu, S., Wang, A., and Au, K. F. (2019) A comparative evaluation of hybrid error correction methods for error-prone long reads. Genome Biology, 20(1), 26.

24. Lima, L., Marchet, C., Caboche, S., Da Silva, C., Istace, B., Aury, J.- M., Touzet, H., and Chikhi, R. (2019) Comparative assessment of long-read error correction software applied to Nanopore RNA-sequencing data. Briefings in bioinformatics,.

